# Epigenetic Activation of Endothelial Smurf1 via EP300-Mediated H3K27ac Disrupts BMPR2 Signaling in Pulmonary Arterial Hypertension

**DOI:** 10.1101/2025.10.30.685664

**Authors:** Celine Leppert, Maria T Ochoa, Joseph D Etlinger, Malik Bisserier

## Abstract

**Background:** Pulmonary arterial hypertension (PAH) is a progressive vascular disease characterized by pulmonary endothelial dysfunction, vascular remodeling, and right ventricular failure. Despite recent advances, the underlying molecular mechanisms remain incompletely understood, and curative treatments are still lacking. Loss of BMPR2 signaling is a hallmark of PAH pathogenesis, yet the mechanisms leading to BMPR2 destabilization are not fully defined. Smad ubiquitination regulatory factor 1 (Smurf1), an E3 ubiquitin ligase, has been implicated in the degradation of BMPR2 and downstream signaling proteins; however, the regulation of Smurf1 and its therapeutic potential remain largely unexplored.

**Methods:** We assessed Smurf1 expression in lung tissues from PAH patients, experimental animal models, and pulmonary artery endothelial cells (PAECs). Epigenetic regulation of Smurf1 was examined using ChIP-qPCR, EP300 gain- and loss-of-function studies, and pharmacological inhibition with the selective EP300 inhibitor A485. Functional consequences of Smurf1 inhibition were evaluated in vitro using transcriptomic profiling, proliferation assays, and BMPR2-Smad signaling analyses. In vivo therapeutic efficacy was tested in monocrotaline (MCT)-induced PAH rats treated with the selective Smurf1 inhibitor Smurf1-IN-A01.

**Results:** Smurf1 expression was elevated in PAH patient lungs, experimental models, and PAECs. Increased histone H3K27 acetylation and EP300 occupancy at the Smurf1 promoter implicated epigenetic activation. EP300 inhibition reduced H3K27ac enrichment, suppressed Smurf1 expression, and restored BMPR2 signaling. Smurf1 blockade with Smurf1-IN-A01 reprogrammed the transcriptomic landscape of PAH-PAECs, downregulating inflammatory, fibrotic, and angiogenic gene networks. Functionally, Smurf1 inhibition decreased pathological endothelial proliferation and enhanced BMPR2-mediated Smad1/5/9 activation. In vivo, Smurf1- IN-A01 improved pulmonary hemodynamics, reduced vascular remodeling and fibrosis, and restored BMPR2 signaling in MCT-PAH rats.

**Conclusions:** Our study identifies Smurf1 as an epigenetically regulated driver of endothelial dysfunction and vascular remodeling in PAH. Targeting Smurf1 restores BMPR2 signaling, reprograms pathogenic processes, and offers a novel therapeutic strategy for PAH.

## INTRODUCTION

Pulmonary arterial hypertension (PAH) is a progressive and multifactorial disease marked by elevated pulmonary arterial pressure, increased pulmonary vascular resistance, and right ventricular failure ^1, 2^. Despite therapeutic advances over the past decade, PAH remains a life-threatening condition with limited treatment options, as current therapies primarily offer modest hemodynamic improvements^3, 4^. A hallmark of PAH is the extensive remodeling of distal pulmonary arteries, driven by the aberrant proliferation, migration, and apoptosis resistance of pulmonary artery endothelial cells (PAECs) and smooth muscle cells (PASMCs)^5^. Among these, endothelial dysfunction is increasingly recognized as an early and pivotal event in PAH pathogenesis^6^, contributing to a pro-proliferative and anti-apoptotic vascular environment by precipitating a cascade of maladaptive vascular responses ^2, 7, 8^. Disruption of endothelial homeostasis leads to an imbalance in vasoactive mediators, favoring vasoconstriction, inflammation, and thrombosis. These alterations promote the proliferation of PASMC and pulmonary artery fibroblasts, contributing to vascular remodeling and increased pulmonary vascular resistance. Furthermore, endothelial dysfunction is associated with processes such as endothelial-to-mesenchymal transition, resistance to apoptosis, and pro-thrombotic states, all key features of PAH pathogenesis and progression. Elucidating the mechanisms underlying endothelial dysfunction is essential for developing targeted therapies to restore vascular homeostasis in PAH^9, 10^. The loss of bone morphogenetic protein receptor type 2 (BMPR2) signaling has emerged as a central driver of endothelial dysfunction in the pulmonary vasculature.

BMPR2 is a member of the transforming growth factor-beta (TGF-β) superfamily. BMPR2 signaling in endothelial cells regulates critical processes, including cell proliferation, differentiation, and apoptosis ^11, 12^. Loss-of-function mutations and reduced BMPR2 expression have been commonly identified in PAH patients, correlating with disease severity and poor clinical outcomes^13–15^. Experimental models with endothelial-specific deletion of BMPR2 signaling recapitulate key features of human PAH, including spontaneous pulmonary vascular remodeling and elevated pulmonary pressures, underscoring the critical role of endothelial BMPR2 signaling in vascular integrity ^6, 16, 17^. While genetic mutations account for some cases of BMPR2 deficiency, emerging evidence highlights the significance of post-translational mechanisms in modulating BMPR2 expression and function^18^. Smad ubiquitination regulatory factor 1 (Smurf1), an E3 ubiquitin ligase, has been implicated in the ubiquitination, proteasomal, as well as lysosomal degradation of BMPR2 and its downstream effectors, Smad1/5 ^19^. Elevated Smurf1 expression has been observed in the pulmonary vasculature of PAH models, leading to impaired BMPR2 signaling ^19^. However, the upstream regulatory mechanisms governing Smurf1 expression in PAH remain poorly understood. Recent studies, including our own, suggest that epigenetic modifications, particularly histone acetylation, play a crucial role in the transcriptional regulation of genes involved in PAH ^20–22^. Alterations in chromatin structure and histone modifications can lead to aberrant gene expression profiles that drive disease progression^23, 24^. Given this context, we hypothesize that Smurf1 upregulation in PAH is mediated through epigenetic mechanisms, contributing to the destabilization of BMPR2 signaling and subsequent endothelial dysfunction.

In this study, we aim to: (1) elucidate the epigenetic mechanisms underlying Smurf1 upregulation in PAH; (2) investigate how Smurf1 modulates BMPR2 expression and downstream Smad signaling in pulmonary endothelial cells; and (3) assess the therapeutic potential of a selective Smurf1 inhibitor, Smurf1-IN-A01, in restoring endothelial function and attenuating pulmonary vascular remodeling. Furthermore, we explore the translational implications of the Smurf1-BMPR2 axis as a novel therapeutic target in the management of PAH.

## MATERIAL AND METHODS

### Human lung tissues

Human lung tissue samples were obtained from the Pulmonary Hypertension Breakthrough Initiative (PHBI), a collaborative biorepository that collects and distributes well-characterized specimens from patients with PAH. Lung tissues were collected from patients with idiopathic PAH (IPAH) at the time of lung transplantation, as well as from non-diseased donor lungs deemed unsuitable for transplantation, serving as healthy controls. All protocols for human tissue procurement were approved by the Institutional Review Boards at New York Medical College. All samples were de-identified prior to distribution. Upon receipt, lung tissues were snap-frozen in liquid nitrogen for RNA and protein extraction. Tissue handling, storage, and processing were conducted according to standardized protocols to ensure sample integrity and comparability across experimental conditions.

### Cell culture

Human pulmonary artery endothelial cells were obtained from PHBI, a multi-center consortium dedicated to collecting and distributing tissue specimens from patients with PAH and control donors. Upon receipt, PAECs were cultured in EC complete medium (Sciencell, Cat# 1001), supplemented with endothelial cell growth Supplement (Sciencell, Cat. #1052), and maintained at 37°C in a humidified atmosphere containing 5% CO. Cells were grown to 80-90% confluence and subcultured at a density of at least 5.0 × 10 cells per 100 mm dish. Only cells between passages 2 and 6 were used for experiments to ensure phenotypic consistency. All cell lines tested negative for mycoplasma, bacteria, yeast, fungi, HIV-1, hepatitis B virus, and hepatitis C virus.

### Reagents and constructs

We used Smurf1-IN-A01 (A01), a potent and selective inhibitor of the E3 ubiquitin ligase Smurf1, purchased from MedChemExpress (MedChemExpress, Cat# HY-110195). A01 has been shown to block Smurf1-mediated ubiquitination and degradation of Smad1/5 proteins, thereby enhancing downstream BMP signaling ^25^. To target epigenetic regulators, we employed A-485 (MedChemExpress, Cat# HY-107455), a selective catalytic inhibitor of the histone acetyltransferases p300 and CBP. A-485 binds competitively to the acetyl-CoA site of p300/CBP, suppressing histone acetylation at lysine 27 of histone H3, a mark associated with transcriptional activation^26^. For gene silencing studies, a lentiviral shRNA construct targeting human *SMURF1* was obtained from Horizon Discovery and cloned into the pLKO.1-puro vector, enabling stable knockdown in PAECs. For *BMPR2* overexpression, we used the pLX304-BMPR2 lentiviral construct, which carries a blasticidin resistance marker and is optimized for high-level gene expression in mammalian systems. Detailed lentiviral construct information, including catalog number and provider, is provided in **Supplementary Table 1**.

### Transient overexpression and siRNA-mediated gene silencing

For transient overexpression studies, human *EP300* (Horizon Discovery, Cat# MHS1010- 202699710) and *SMURF1* expression constructs (Horizon Discovery, Cat# MHS1010- 202696309) were transfected into FD-PAECs using Lipofectamine™ 2000 (Thermo Fisher, Cat#11668019) according to the manufacturer’s protocol. Cells were transfected with 2 µg plasmid DNA per well in 6-well plates, and the empty pCMV·SPORT-βgal vector (Thermo Fisher, Cat#10586014) was used as a negative control. Cells were harvested for total RNA and protein extraction 48 hrs post-transfection. For gene silencing, PAH-PAECs were treated with Accell siRNA-targeting *EP300* (Horizon Discovery, Cat# A-003486-19-0005) or non-targeting siRNA as a control vector (Horizon Discovery, Cat# D-001910-01-05). siRNA delivery was performed in serum-free media, following the manufacturer’s instructions. Cells were processed for downstream assays, including RNA and protein analysis, after 72 hrs of incubation with siRNA.

### Animal Models

All animal experiments were conducted in accordance with the National Institutes of Health Guide for the Care and Use of Laboratory Animals and were approved by the Institutional Animal Care and Use Committee (IACUC) of New York Medical College. Male C57BL/6J mice (8–10 weeks old; Jackson Laboratory) were subjected to the SuHx protocols, consistent with established protocols ^27–29^. Briefly, adult mice received weekly subcutaneous injections of SU5416 (20 mg/kg; MedChemExpress), a selective VEGFR 2 inhibitor, for three consecutive weeks. Immediately after each injection, mice were placed in a normobaric hypoxia chamber (10% O ; ProOx 360, BioSpherix Ltd). After the final hypoxic exposure, animals were returned to normal oxygen conditions for a 7 day recovery in normoxia. PAH induction in mice was confirmed by measuring right ventricular systolic pressure (RVSP) and assessing right ventricular hypertrophy using the Fulton index (RV/[LV+S]). Male Sprague-Dawley rats (200– 225 g; Charles River Laboratories) were used for the monocrotaline (MCT)-induced model of PAH. Rats received a single subcutaneous injection of monocrotaline (60 mg/kg; Sigma-Aldrich) to induce PAH. After 21 days, rats were randomized to receive either vehicle control or Smurf1- IN-A01 (20 mg/kg) administered intraperitoneally every other day for an additional 14 days. At day 35, RVSP and mean pulmonary artery pressure (mPAP) were assessed via right heart catheterization as described below. Following hemodynamic assessment, lung and heart tissues were harvested for histological and molecular analyses. Both the SuHx and MCT samples analyzed in **Figure 1** of this study were obtained from experimental cohorts validated and reported in previously published studies {34078089}, in which PH was confirmed by elevated RVSP, increased Fulton index, distal pulmonary artery remodeling, and elevated PAP in MCT rats.

**Figure 1.**
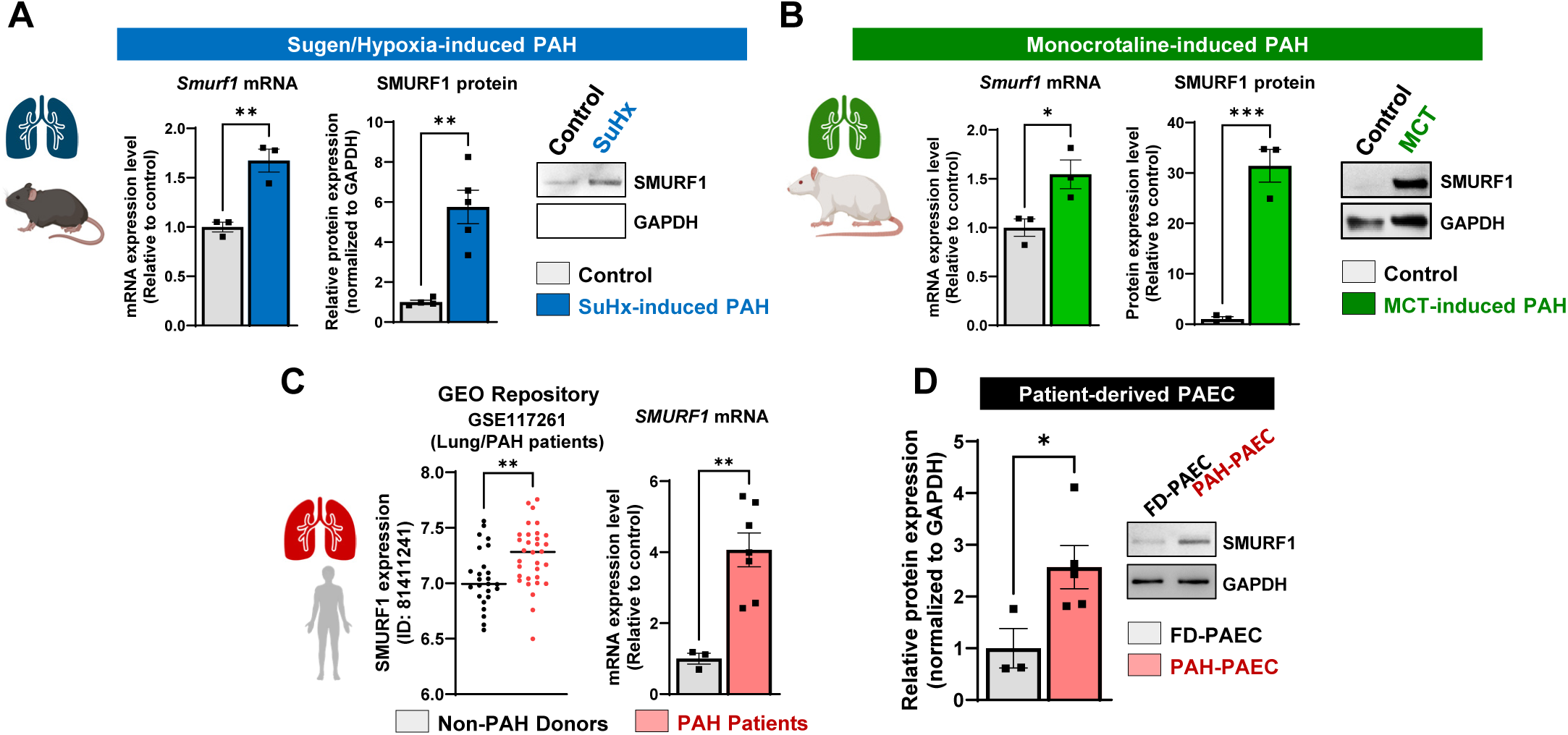
Smurf1 expression is upregulated in experimental and human pulmonary arterial hypertension. **(A)** SMURF1 expression measured by quantitative PCR and western blot analysis in lung tissues from SU5416/hypoxia (SuHx)-exposed mice, compared to normoxic controls (n=3-5). **(B)** SMURF1 levels in lungs from monocrotaline (MCT)-treated rats assessed by quantitative PCR and western blot analysis (n=3). **(C)** Analysis of public RNA-seq dataset GSE117261 (left panel) for *SMURF1* transcript levels in PAH patient lungs compared to non-PAH controls (n=28-32). *SMURF1* transcript levels were measured by qRT-PCR (right panel) in lung tissues from non-PAH and IPAH patients (n=3-7). **(D)** SMURF1 protein expression was assessed in FD and PAH-PAECs by immunoblotting. Data are presented as mean ± SEM. *p<0.05; **p<0.01, ***p<0.001.

### Hemodynamic Measurements via Right Heart Catheterization

RVSP and mPAP were measured in anesthetized rats to assess pulmonary hemodynamics. Animals were anesthetized with 2-4% isoflurane and intubated via tracheotomy. Mechanical ventilation was maintained with 1-2% isoflurane in oxygen. A thoracotomy was performed to expose the heart and great vessels. A 1F Millar Mikro-Tip® pressure-volume catheter (PVR- 1030; Millar Instruments) was inserted directly into the right ventricle through the free wall to record RVSP. For mPAP measurements, the catheter was advanced through the right ventricular outflow tract into the main pulmonary artery under continuous pressure waveform monitoring. The catheter was connected to a PowerLab data acquisition system, and pressure waveforms were recorded and analyzed using LabChart software.

### Lentivirus preparation

Lentiviral particles were produced using the second-generation packaging system. HEK293T cells were co-transfected with the transfer plasmid (e.g., pLKO.1-shRNA or pLenti- overexpression construct), packaging plasmid psPAX2 (Addgene, #12260), and envelope plasmid pMD2.G (Addgene, #12259) using Lipofectamine 2000 (Invitrogen, #11668019), following the manufacturer’s protocol. Approximately 12 hours post-transfection, the medium was replaced with fresh DMEM supplemented with 10% fetal bovine serum (FBS) and without antibiotics. After 48 hours, the viral supernatant was collected and filtered through a 0.2 μm polyethersulfone filter to remove cellular debris. This protocol was adapted from and is consistent with methods previously described in our earlier studies ^30^. The efficiency of gene knockdown or overexpression was validated by immunoblotting and quantitative reverse transcription PCR (RT-qPCR) to assess changes in protein and mRNA expression levels, respectively.

### In vitro cell proliferation

PAECs were cultured for 48 hours under various conditions: treatment with sh*SMURF1* (a lentiviral-mediated knockdown of SMURF1), Smurf1-IN-A01 (Smurf1 inhibitor), and lentiviral-mediated overexpression of *BMPR2*. Corresponding control treatments were applied in parallel. Following treatment, cells were fixed with 4% paraformaldehyde for 15 minutes and rinsed with phosphate-buffered saline (PBS). Cell proliferation was assessed via immunofluorescence staining for Ki67 (Thermo Fisher, Cat# MA5-14520) and proliferating cell nuclear antigen (PCNA, Thermo Fisher, Cat# PA1-38424), established markers of cellular proliferation. Nuclei were counterstained with DAPI (Thermo Fisher, Cat# 62248) to determine the total cell count. Proliferation levels were quantified by calculating the ratio of Ki67- or PCNA-positive cells to the total number of DAPI-stained nuclei per field. Each experimental condition was performed in triplicate, with a minimum of 3,000 cells counted per condition to ensure statistical robustness. Additionally, the MTT assay (Sigma-Aldrich, Cat# M2128) was employed to evaluate cell viability and proliferation. After 48 hours of treatment, 10 µL of MTT reagent (5 mg/mL) was added to each well containing 100 µL of culture medium, and the cells were incubated at 37°C for 4 hours. Subsequently, 100 µL of solubilization solution (e.g., DMSO) was added to dissolve the formazan crystals, and the absorbance was measured at 570 nm using a microplate reader. The absorbance values directly correlate with the number of metabolically active cells, providing a quantitative measure of cell viability and proliferation.

### Hematoxylin & Eosin and Masson’s trichrome staining

Lung tissues were fixed in 4% formaldehyde, embedded in paraffin, and sectioned at 5 μm thickness. Sections underwent deparaffinization in xylene and rehydration through graded alcohols. Hematoxylin and eosin (H&E) staining was performed to assess vascular remodeling, while Masson’s trichrome staining was used to evaluate collagen deposition and fibrosis. For Masson’s trichrome staining, nuclei were stained with Weigert’s iron hematoxylin, cytoplasm with Biebrich scarlet-acid fuchsin, and collagen fibers with aniline blue, following established protocols. Stained sections were examined under light microscopy. Quantitative analysis of medial thickness and collagen deposition was conducted using ImageJ software, employing color deconvolution and thresholding techniques to isolate and measure the areas of interest, as previously described in our earlier studies ^21, 31, 32^.

### Western Blot

Total protein was extracted from cell or tissue samples using RIPA lysis buffer (Invitrogen, Cat# 89901), supplemented with protease and phosphatase inhibitor cocktail (Thermo Fisher, Cat# A32959). Lysates were clarified by centrifugation at 15,000 × g for 20 minutes at 4°C, and protein concentrations were determined using the bicinchoninic acid assay (Thermo Fisher, Cat# A55865). Equal amounts of protein (typically 20–40 µg) were resolved on SDS-polyacrylamide gels and transferred onto nitrocellulose membranes using the Trans-Blot Turbo Transfer System (Bio-Rad). Membranes were blocked for 1 hour at room temperature with either 5% non-fat dry milk or 5% BSA in TBS-Tween (0.1%) and incubated overnight at 4°C with primary antibodies (listed in **Supplemental Table 1**). Following washes, membranes were incubated with HRP-conjugated secondary antibodies (Cell Signaling Technology) for 1 hour at room temperature. Signal detection was performed using enhanced chemiluminescence (Thermo Fisher, Cat# 34075), and bands were visualized with a ChemiDoc imaging system (Bio-Rad). Densitometric quantification of protein bands was conducted using ImageJ software, with target protein levels normalized to the corresponding loading control.

### Quantitative real-time PCR

Total RNA was isolated from right ventricular (RV) tissues, lung specimens, and PAECs using TRIzol reagent (Invitrogen, Cat# 15596018), in accordance with the manufacturer’s instructions. RNA purity and concentration were assessed via spectrophotometric analysis. Complementary DNA was synthesized from 1 µg of total RNA using the qScript cDNA Synthesis Kit (Quantabio, Cat# 95217), which employs a combination of oligo(dT) and random primers to ensure unbiased reverse transcription across transcript species. Quantitative real-time PCR was performed in technical triplicate on the QuantStudio 6 Pro Real-Time PCR System (Applied Biosystems) using SYBR Green-based detection (Applied Biosystems, Cat# A46112). Amplification reactions incorporated gene-specific primers (sequences listed in **Supplemental Table 1**). Specificity of amplification products was verified by melting curve analysis. Relative transcript levels were determined by the comparative threshold cycle (ΔΔCt) method, with 18S rRNA serving as the endogenous normalization control.

### mRNA Sequencing and Transcriptome Data Analysis

Total RNA was extracted from PAECs using RNeasy Mini Kits (Qiagen, Cat# 74106) according to the manufacturer’s protocol. RNA purity and concentration were assessed with a NanoDrop spectrophotometer, and RNA integrity was evaluated using an Agilent 2100 Bioanalyzer. Samples exhibiting RNA integrity numbers (RIN) >7.0, OD260/280 ≥2.0, and OD260/230 ≥2.0 were considered suitable for sequencing. RNA quality was further confirmed by electrophoresis on 1% agarose gels to exclude degradation and contamination. RNA sequencing was performed by Novogene (Sacramento, CA) using the Illumina NovaSeq 6000 platform with 250–300 bp insert size cDNA libraries and a 150 bp paired-end read configuration. For each condition, three biological replicates were sequenced. Differential gene expression analysis between groups was performed using the DESeq2 R package, which models count data using a negative binomial distribution. Resulting p-values were corrected for multiple testing using the Benjamini-Hochberg method to control the false discovery rate (FDR). Genes were considered differentially expressed if they met an adjusted p-value (FDR) threshold of <0.05 and an absolute log2 fold- change ≥1.5. Log2 fold changes were plotted against -log10 adjusted p-values in volcano plots to visualize gene-level significance and magnitude of expression differences. Prior to clustering and heatmap generation, gene counts were normalized using the logCPM transformation. The top 2,500 most variable genes were selected, and Z-score standardized for hierarchical clustering. Heatmaps were generated using Clustergrammer (http://amp.pharm.mssm.edu/clustergrammer/), an interactive web-based tool for exploring high-dimensional transcriptomic data.

### Chromatin immunoprecipitation followed by quantitative PCR (ChIP–qPCR)

ChIP assays were performed using the EZ-Magna ChIP™ A/G Chromatin Immunoprecipitation Kit (Millipore, Cat# 17-10086) following the manufacturer’s protocol. Briefly, PAECs were cross-linked with 1% formaldehyde for 10 minutes at room temperature to preserve protein-DNA interactions, and the reaction was quenched with 125 mM glycine. Cells were then washed with cold PBS, harvested, and lysed sequentially using Cell Lysis Buffer and Nuclear Lysis Buffer provided in the kit. Chromatin was sheared to an average length of 200–500 bp by sonication. For immunoprecipitation, 5 µg of anti-H3K27ac antibody (Cell Signaling Technology, Cat# 8173S) was incubated with the sheared chromatin overnight at 4°C with rotation. Protein A/G magnetic beads were then added to capture the antibody-chromatin complexes. After extensive washing to reduce nonspecific binding, the complexes were eluted, and cross-links were reversed by heating at 65°C for 4 hours. DNA was purified using spin columns included in the kit. Quantitative PCR was performed using Fast SYBR Green Master Mix (Applied Biosystems, Cat# A46112) with primers specific against the SMURF1 promoter (primer sequences listed in **Supplemental Table 1**). Enrichment of H3K27ac at the SMURF1 promoter was calculated as a percentage of input DNA and normalized to IgG controls to confirm specificity. Fold changes in enrichment between experimental groups were expressed relative to control samples.

### Statistical Analysis

All statistical analyses were performed using GraphPad Prism (version 10.3.0, GraphPad Software, San Diego, CA, USA), and data were presented as mean ± standard error of the mean (SEM). The Shapiro-Wilk test was first conducted to assess data normality. For comparisons between two independent groups, an unpaired t-test was used for normally distributed data, whereas the Mann-Whitney U test was applied for non-normally distributed data. When comparing more than two groups, a one-way ANOVA was performed if data met normality assumptions, followed by Tukey’s multiple comparisons test for pairwise comparisons or Dunnett’s test when comparing against a control group. For non-normally distributed data, the Kruskal-Wallis test was used, followed by Dunn’s post-hoc test for multiple comparisons. To assess the effects of two independent variables, a two-way ANOVA was performed, with post-hoc tests conducted when significant interactions were detected. Statistical significance was denoted as p < 0.05 (*), p < 0.01 (**), p < 0.001 (***). All experiments were conducted with a minimum of three biological replicates.

## RESULTS

### Smurf1 upregulation in preclinical models and human pulmonary arterial hypertension

To elucidate the potential involvement of Smurf1 in the pathobiology of PAH, we systematically examined its expression across both rodent models and human-derived tissues. In the SU5416/hypoxia (SuHx) model of experimental PAH, which replicates key features of severe human disease, including endothelial dysfunction and vascular obliteration^33^, *Smurf1* mRNA and protein levels were significantly increased relative to normoxic controls in mice, supporting its potential role in pulmonary vascular remodeling (**Figure 1A**). Consistent with these findings, *Smurf1* expression was also markedly upregulated at the transcript and protein levels in MCT-treated rats compared with untreated controls **(Figure 1B**), supporting a conserved role for Smurf1 across distinct preclinical models of PAH. Both the SuHx and MCT samples used in this study were obtained from animal models that had been validated and described in previously published studies {34078089}. We next examined Smurf1 expression in human PAH by analyzing the publicly available RNA-seq dataset GSE117261, which confirmed elevated *SMURF1* transcript levels in lung tissues from PAH patients compared with non-PAH donors (**Figure 1C, left panel**). Independent validation by quantitative RT-PCR demonstrated a significant increase in *SMURF1* mRNA expression in lung samples from IPAH patients relative to controls (**Figure 1C, right panel**). To further extend these findings, we assessed SMURF1 expression in pulmonary artery endothelial cells isolated from IPAH patients (PAH-PAECs). Both qPCR and immunoblot analyses confirmed that SMURF1 was significantly upregulated at the mRNA and protein levels in PAH-PAECs compared with donor-derived endothelial cells (FD-PAECs) (**Figure 1D**). Together, these results demonstrate that SMURF1 expression is consistently elevated across experimental models and clinical PAH samples, implicating Smurf1 as a potential mediator of pulmonary vascular remodeling.

### EP300-driven H3K27 acetylation epigenetically promotes Smurf1 expression in PAH

Epigenetic modifications have emerged as critical regulators of gene expression in PAH, as they control the transcriptional programs that drive vascular remodeling and endothelial dysfunction^34,35^. Among these, histone H3 lysine 27 acetylation (H3K27ac) is a well-characterized epigenetic mark associated with active enhancer and promoter regions, facilitating an open chromatin state and promoting transcriptional activation^36^ ^24^. Previous studies have demonstrated that H3K27ac levels are significantly elevated in the lungs of PAH patients and fibroblasts derived from PAH patients^37, 38^, suggesting a broader role for histone acetylation in the pathogenesis of vascular remodeling and fibrosis in PAH. To investigate whether Smurf1 expression is regulated through epigenetic mechanisms, we first analyzed publicly available ENCODE ChIP-seq datasets and identified marked enrichment of H3K27ac across the Smurf1 promoter, consistent with active transcriptional regulation (**Figure 2A**). In silico analysis further revealed enriched binding sites for key transcriptional coactivators, including EP300 (p300) and CCAAT/enhancer-binding protein beta (CEBPB) (**Figure 2A**). EP300, a histone acetyltransferase (HAT), catalyzes the acetylation of lysine residues on histone tails, facilitating an open chromatin structure that favors transcriptional activation. To interrogate the functional role of EP300 in regulating SMURF1 expression, we employed both gain and loss-of-function strategies in FD-PAECs and PAH-PAECs, respectively. In FD-PAECs, *EP300* overexpression led to an increase in *SMURF1* mRNA levels **(Figure 2B)**, accompanied by elevated global H3K27 acetylation and enhanced SMURF1 protein abundance **(Figure 2C)**. In contrast, *EP300* depletion in PAH-PAECs significantly reduced both *EP300* and *SMURF1* transcript levels **(Figure 2D)**. Western blot analysis confirmed decreased H3K27ac and SMURF1 protein levels in *EP300*-depleted PAH-PAECs **(Figure 2E)**.

**Figure 2.**
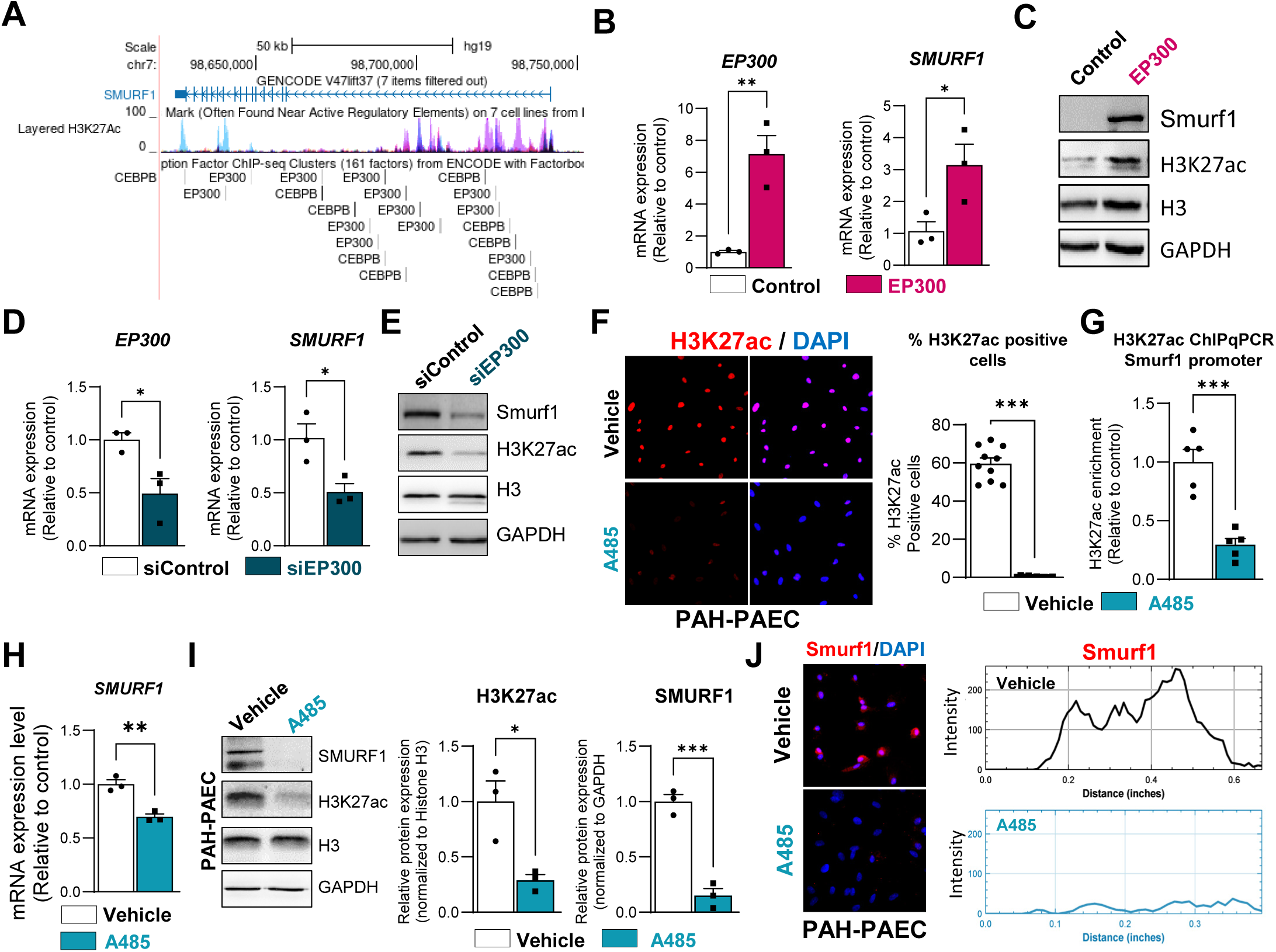
EP300-dependent histone acetylation promotes Smurf1 transcription in PAH endothelial cells. **(A)** ChIP-seq data from ENCODE reveal H3K27ac enrichment at the Smurf1 promoter region, with predicted EP300 and CEBPB binding motifs. **(B)** Quantitative PCR analysis of *EP300* and *SMURF1* mRNA expression in FD-PAECs following *EP300* overexpression, n=4. **(C)** Immunoblot against global H3K27ac and SMURF1 protein levels in FD-PAECs, 48 hrs after EP300 overexpression. GAPDH and H3 were used as loading controls, respectively. **(D)** Quantitative PCR analysis of *EP300* and *SMURF1* mRNA expression in *EP300*-depleted PAH-PAECs, n=4. **(E)** Immunoblot analysis of global H3K27ac and SMURF1 protein levels in PAH-PAECs, 72 hrs after after siRNA delivery. **(F)** PAECs derived from PAH patients were treated with A485 (2 µM, 48 hrs), and global H3K27ac levels were assessed by immunofluorescence, n=3. **(G)** ChIP-qPCR quantifies H3K27ac enrichment at the Smurf1 promoter following A485 treatment, n=4. **(H-I)** *SMURF1* mRNA and protein levels were measured by qPCR and Western blot, respectively, after EP300 inhibition, n=3. **(J)** Smurf1 expression was assessed by immunofluorescence in control and A485-treated PAH-PAECs, n=3. Data are presented as mean ± SEM. *p<0.05; **p<0.01, ***p<0.001.

To complement these genetic approaches and further dissect the epigenetic dependency of SMURF1 expression on EP300 catalytic activity, we employed the selective EP300/CBP inhibitor A485 in PAH-derived PAECs. Treatment of PAH-derived PAECs with A485 significantly reduced global H3K27ac levels, as shown by immunofluorescence staining (**Figure 2F**). Indeed, quantitative analysis confirmed a decreased proportion of H3K27ac-positive cells following EP300 inhibition, demonstrating effective suppression of histone acetylation using A485. To specifically assess changes at the Smurf1 promoter, we performed ChIP-qPCR and found a marked decrease in H3K27ac enrichment within the *SMURF1* promoter region upon A485 treatment (**Figure 2G**). Consistent with these chromatin-level changes, *SMURF1* mRNA levels were significantly downregulated following EP300 inhibition (**Figure 2H**). Western blot analysis confirmed reductions in Smurf1 protein expression and global H3K27ac levels (**Figure 2I**). Consistently, immunofluorescence staining further validated the decrease in Smurf1 protein following A485 treatment (**Figure 2J**). Together, these findings identify EP300-mediated histone acetylation as a key epigenetic mechanism promoting Smurf1 overexpression in PAH endothelial cells. Our results also highlight EP300 as a potential therapeutic target to normalize aberrant gene expression programs driving pulmonary vascular remodeling and disease progression.

### Smurf1-mediated BMPR2 degradation induces endothelial dysfunction in PAH

Previous studies have suggested that Smurf1 targets BMPR2 and drives its ubiquitin-mediated proteasomal and lysosomal degradation ^19, 39^, contributing to the impairment of BMP signaling pathways ^40^. To investigate this potential interaction, we first used the STRING protein-protein interaction database, which confirmed a direct association between Smurf1 and BMPR2 (**Figure 3A)**. To validate these findings experimentally, we conducted loss-of-function studies in PAH-PAECs using a short-hairpin RNA approach specifically targeting *SMURF1*. Efficient knockdown of *SMURF1* was confirmed at both the mRNA and protein levels compared to control-infected cells **(Figure 3B)**. Notably, *SMURF1* depletion significantly increased BMPR2 protein expression and enhanced phosphorylation of the downstream Smad1/5/9 signaling molecules **(Figure 3B)**, indicating a restoration of BMP signaling activity. To complement the genetic approach, we employed a pharmacological strategy using Smurf1-IN-A01, a recently developed small-molecule inhibitor that selectively targets the E3 ubiquitin ligase activity of Smurf1. Smurf1-IN-A01 functions by disrupting the interaction between Smurf1 and its ubiquitin-conjugating enzymes, thereby preventing the degradation of Smurf1 substrates such as BMPR2. Importantly, Smurf1-IN-A01 exhibits high specificity for Smurf1 over related HECT domain-containing E3 ligases, minimizing potential off-target effects. Treatment of PAECs with Smurf1-IN-A01 recapitulated the effects observed with genetic silencing, leading to increased BMPR2 expression and enhanced phosphorylation of Smad1/5/9 **(Figure 3C)**. Collectively, these data demonstrated that Smurf1 acts as a negative regulator of BMPR2 signaling in PAECs by promoting its degradation. Furthermore, they support the rationale for targeting Smurf1 as a therapeutic strategy to restore BMP signaling and counteract endothelial dysfunction in PAH.

**Figure 3.**
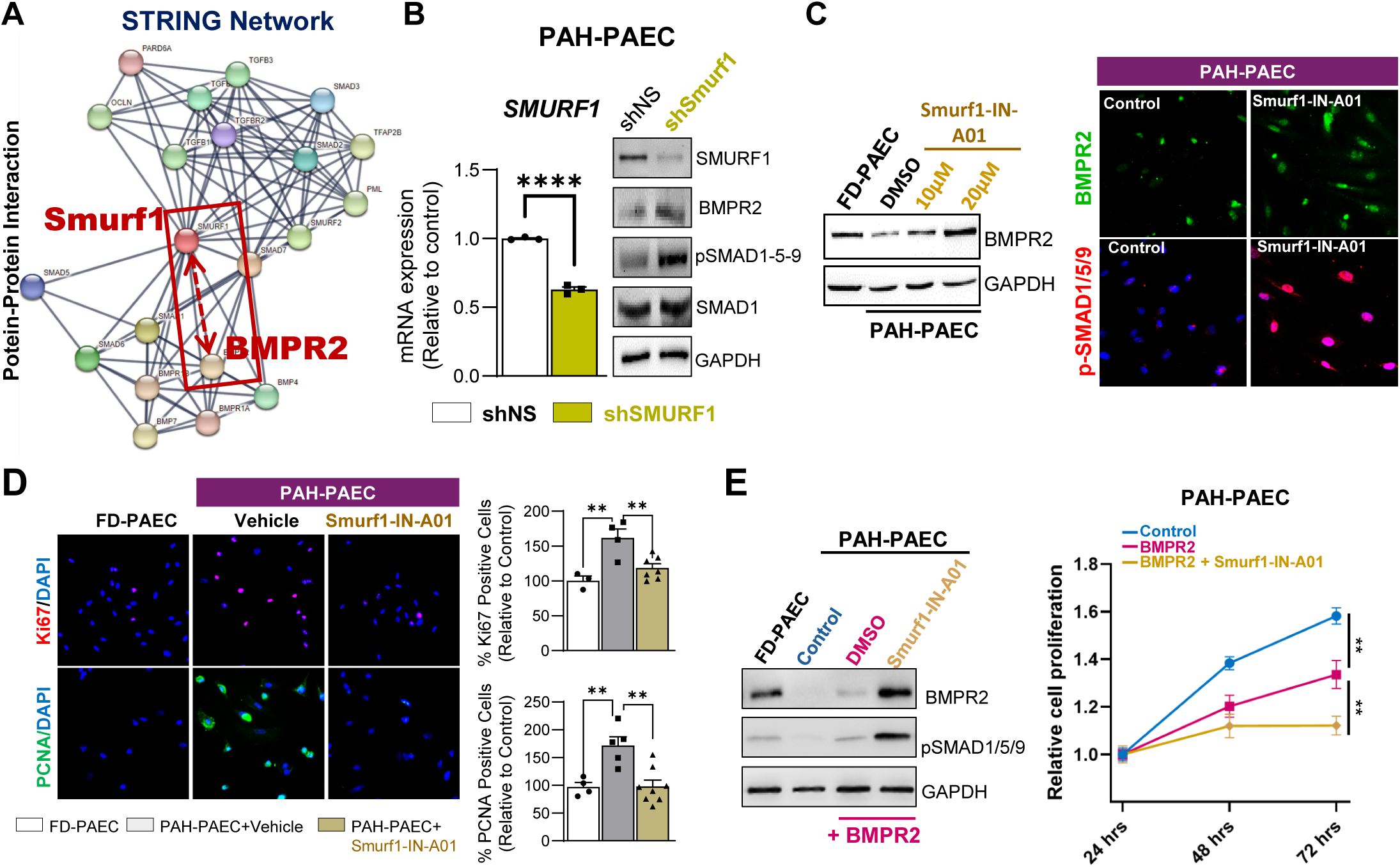
SMURF1 inhibition restores BMPR2 signaling and limits endothelial proliferation in PAH-derived PAECs. **(A)** STRING database analysis identifies a functional interaction between Smurf1 and BMPR2. **(B)** BMPR2 protein levels and Smad1/5/9 phosphorylation in FD-PAECs and PAH-PAECs after *SMURF1* knockdown via lentiviral shRNA, n=3. **(C)** BMPR2 expression and Smad signaling following pharmacologic inhibition of SMURF1 with Smurf1-IN-A01 (20 µM; 72 hrs), as analyzed by immunoblot and immunostaining, n=3. **(D)** Endothelial proliferation was evaluated by immunostaining for Ki67 and PCNA following Smurf1-IN-A01 treatment in FD-PAEC and PAH-PAEC, n=3. **(E)** Combined *BMPR2* overexpression and Smurf1-IN-A01 treatment synergistically upregulate Smad signaling and reduce PAH-PAEC proliferation, n=6. Data are presented as mean ± SEM. *p<0.05; **p<0.01, ***p<0.001.

### Smurf1 blockade reduces aberrant pulmonary endothelial cell proliferation

Excessive proliferation of PAECs is a hallmark feature of PAH, contributing directly to vascular remodeling, luminal obstruction, and progressive elevation of pulmonary vascular resistance. Given our findings that Smurf1 negatively regulates BMPR2 signaling, a pathway known to suppress endothelial proliferation, we next investigated the functional consequences of Smurf1 inhibition in PAH-PAECs. Treatment with the selective Smurf1 inhibitor Smurf1-IN-A01 significantly attenuated the proliferative phenotype of PAH-PAECs, as evidenced by a marked reduction in the number of Ki67- and PCNA-positive cells compared to vehicle-treated controls (**Figure 3D**). These results indicate that pharmacologic blockade of Smurf1 not only restores BMPR2 signaling but also effectively suppresses pathological endothelial proliferation, a key driver of vascular remodeling in PAH. To further define the relationship between Smurf1 inhibition and BMPR2 signaling, we evaluated whether *BMPR2* overexpression alone would be sufficient to rescue signaling defects in PAH-PAECs. We found that BMPR2 overexpression moderately increased receptor expression and promoted Smad1/5/9 phosphorylation; however, it failed to fully restore BMP signaling to levels observed in non-diseased PAECs (**Figure 3E**). Strikingly, simultaneous inhibition of SMURF1 in BMPR2-overexpressing PAECs significantly potentiated BMPR2 protein accumulation and further enhanced Smad1/5/9 phosphorylation compared to *BMPR2* overexpression alone (**Figure 3E, left panel**). This combined approach also led to a more pronounced reduction in endothelial cell proliferation than either intervention alone, suggesting that Smurf1 inhibition synergizes with BMPR2 overexpression to amplify downstream BMP signaling and promote endothelial quiescence (**Figure 3E, right panel**). Overall, our findings provide strong mechanistic evidence that Smurf1 drives pathological endothelial proliferation in PAH and underscore the therapeutic potential of combinatorial strategies targeting both Smurf1 inhibition and BMPR2 restoration to reverse vascular remodeling.

### Smurf1 inhibition reprograms inflammatory and angiogenic signaling in PAH-PAECs

Next, we performed transcriptomic profiling to characterize the global molecular impact of Smurf1 inhibition in PAECs. PAECs derived from PAH patients were treated either with vehicle control or the selective Smurf1 inhibitor Smurf1-IN-A01, and RNA sequencing (RNA-seq) was performed to assess changes in gene expression (**Figure 4A**). Principal component analysis (PCA) revealed a distinct separation between vehicle-treated and Smurf1-IN-A01–treated cells, indicating a robust shift in the global transcriptomic profile following Smurf1 inhibition (**Figure 4B**). Heatmap analysis of biological replicates confirmed reproducible and consistent patterns of differentially regulated genes between treatment groups (**Figure 4C, left panel**). Volcano plot visualization identified 59 significantly upregulated genes and 84 downregulated genes upon Smurf1 inhibition compared to controls (**Figure 4C, right panel**). Gene ontology (GO) enrichment analysis of differentially expressed genes highlighted biological pathways predominantly associated with inflammatory responses, cell migration, tube development, and angiogenesis. Notably, pathways related to tumor necrosis factor (TNF) signaling, angiogenesis, inflammatory cytokine production, and vascular remodeling were significantly enriched following Smurf1 inhibition (**Figure 4D**). To further confirm and extend these findings, we performed enrichment analysis using multiple independent databases, including BioPlanet, ChEA (ChIP-X Enrichment Analysis), MSigDB Hallmark 2020, and GO Biological Processes 2023. These analyses consistently revealed significant enrichment of pathways related to TGF-β signaling regulation of extracellular matrix organization, TNF-α signaling, FOXM1 transcriptional networks, and BRD4-mediated gene regulation (**Figure 4E**). Gene set enrichment analysis (GSEA) further confirmed significant suppression of pro-fibrotic and inflammatory pathways following Smurf1 inhibition, including TGF-β signaling, TNF-α signaling, inflammatory response, and interferon-α response gene sets (**Figure 4F**). Finally, to experimentally validate our RNA-seq findings, we treated PAH-derived PAECs with the selective SMURF1 inhibitor Smurf1-IN-A01 and assessed the expression of canonical inflammatory markers by quantitative PCR. Pharmacological inhibition of SMURF1 significantly reduced *IL1B, IL6, IL8,* and *TNFA* transcript levels relative to vehicle-treated controls (**Figure 4G**), consistent with transcriptomic predictions. To further delineate the direct contribution of SMURF1 to endothelial inflammation, we conducted complementary genetic perturbation studies (**Supplementary** Figure 1). shRNA-mediated silencing of *SMURF1* markedly suppressed *IL6* and *TNFA* expression (**Supplementary** Figure 1A), closely mirroring the effects observed with pharmacological inhibition (**Figure 4G**). Conversely, *SMURF1* overexpression in donor-derived PAECs significantly upregulated *IL6* and *TNFA* mRNA levels (**Supplementary** Figure 1B-C), reinforcing a direct pro-inflammatory role. Collectively, these data establish SMURF1 as a key regulator of inflammatory signaling in the pulmonary endothelium.

**Figure 4.**
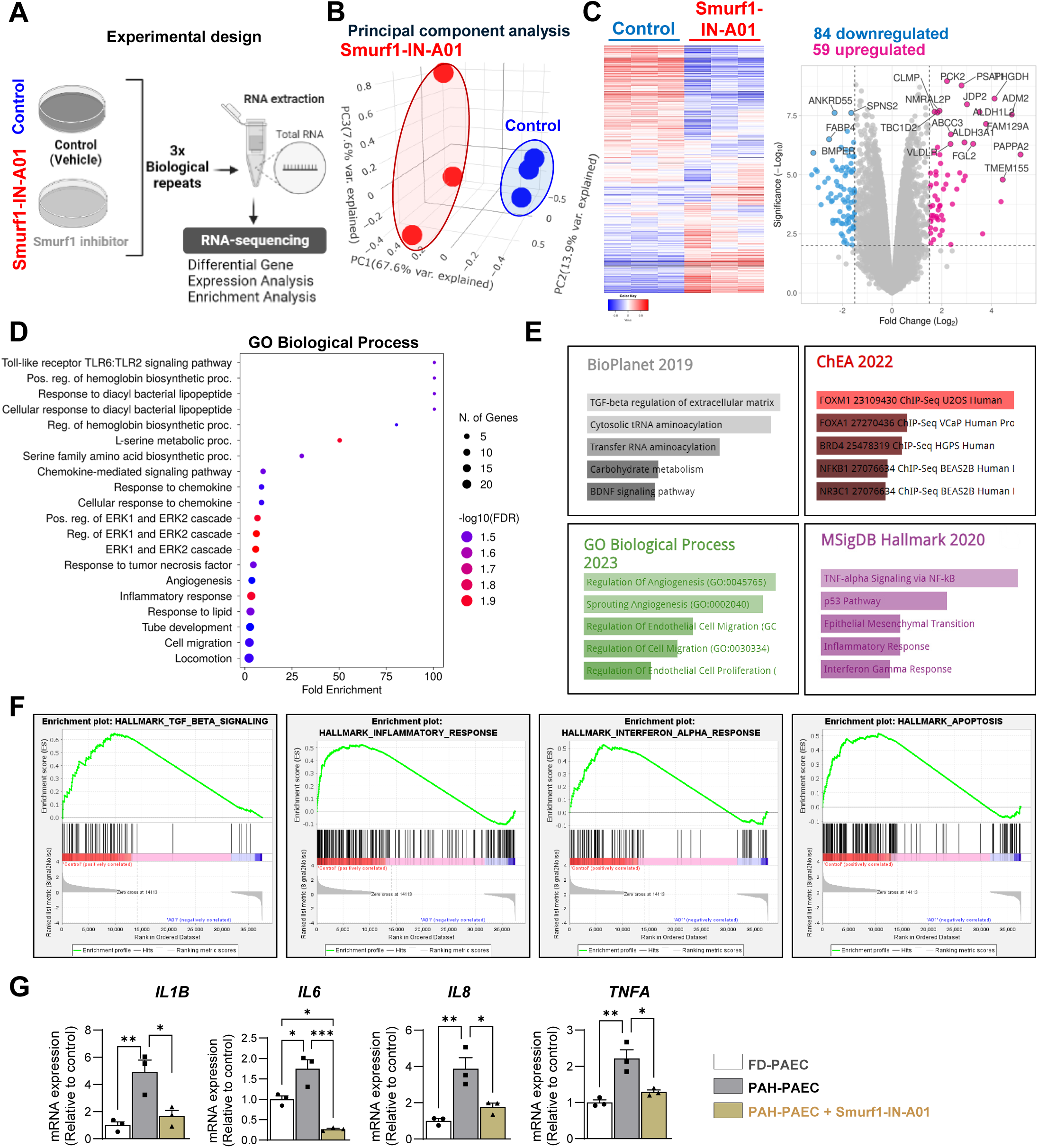
Transcriptomic profiling identifies Smurf1-dependent gene networks in PAH-PAECs. **(A)** RNA-seq workflow for PAH-PAECs treated with vehicle or Smurf1-IN-A01. **(B)** Principal component analysis (PCA) reveals distinct clustering between treated and control groups. **(C)** Heatmap (left) and volcano plot (right) depict global transcriptomic changes following Smurf1 inhibition. **(D)** Gene ontology (GO) enrichment identifies downregulated pathways involved in inflammation, angiogenesis, and ECM remodeling. **(E-F)** Gene Set Enrichment Analysis (GSEA) confirms reduced activation of TGF-β, TNF-α, and interferon-related pathways. **(G)** Inflammatory gene expression (*IL1B, IL6, IL8, TNFA*) was measured by qPCR in Smurf1-IN-A01–treated PAH-PAECs, n=3. Data are presented as mean ± SEM. *p<0.05; **p<0.01, ***p<0.001.

### Targeting Smurf1 improves cardiopulmonary hemodynamics and reverses pulmonary vascular remodeling

Given our in vitro findings that SMURF1 contributes to endothelial dysfunction through suppression of BMPR2 signaling, we next investigated whether pharmacological inhibition of Smurf1 could reverse pulmonary vascular remodeling and improve cardiopulmonary hemodynamics in vivo. To this end, we employed the MCT-induced rat model of PAH, which reproduces cardinal features of the human disease, including elevated pulmonary artery pressures, medial wall thickening, and RV hypertrophy. This inflammatory-driven model is widely used to assess the therapeutic efficacy of candidate compounds targeting pulmonary vascular remodeling. Through this approach, we aimed to determine whether selective inhibition of Smurf1 could restore vascular homeostasis and alleviate disease progression in a clinically relevant experimental context. Treatment of MCT-PAH rats with the selective Smurf1 inhibitor Smurf1-IN-A01 significantly improved pulmonary hemodynamics. RVSP and mPAP were both markedly reduced in Smurf1-IN-A01-treated animals compared to untreated controls (**Figure 5A-B**). Furthermore, Smurf1 inhibition significantly attenuated right ventricular hypertrophy, as reflected by a decreased Fulton index. Histological evaluation of lung tissues using H&E staining revealed a substantial reduction in pulmonary vascular remodeling following Smurf1-IN-A01 treatment, with decreased medial wall thickness of small pulmonary arteries compared to MCT controls (**Figure 5C**). In parallel, Masson’s trichrome staining demonstrated a marked reduction in pulmonary fibrosis in treated rats (**Figure 5D**), findings that were corroborated at the molecular level by a significant decrease in fibrosis-related gene expression, including *Col1a1, Col3a1*, and *Tgfb1* (**Figure 5E**). Moreover, Smurf1 inhibition attenuated inflammation and oxidative stress pathways (**Figure 5F-G**). Smurf1-IN-A01–treated lungs exhibited reduced expression of pro-inflammatory cytokines (*Il6, Tnfa*) and oxidative stress markers (*Nox4, Sod2, Gpx1*), compared to vehicle-treated animals (**Figure 5F-G**). Finally, western blot analysis confirmed that SMURF1 inhibition restored BMPR2 protein expression and enhanced downstream SMAD1/5/9 phosphorylation in lung tissues (**Figure 5H**), suggesting functional reactivation of BMP signaling in vivo. Our data demonstrate that Smurf1 inhibition improves pulmonary hemodynamics, reduces vascular remodeling and fibrosis, and restores BMPR2 signaling, supporting Smurf1 as a promising therapeutic target to reverse disease progression in PAH.

**Figure 5.**
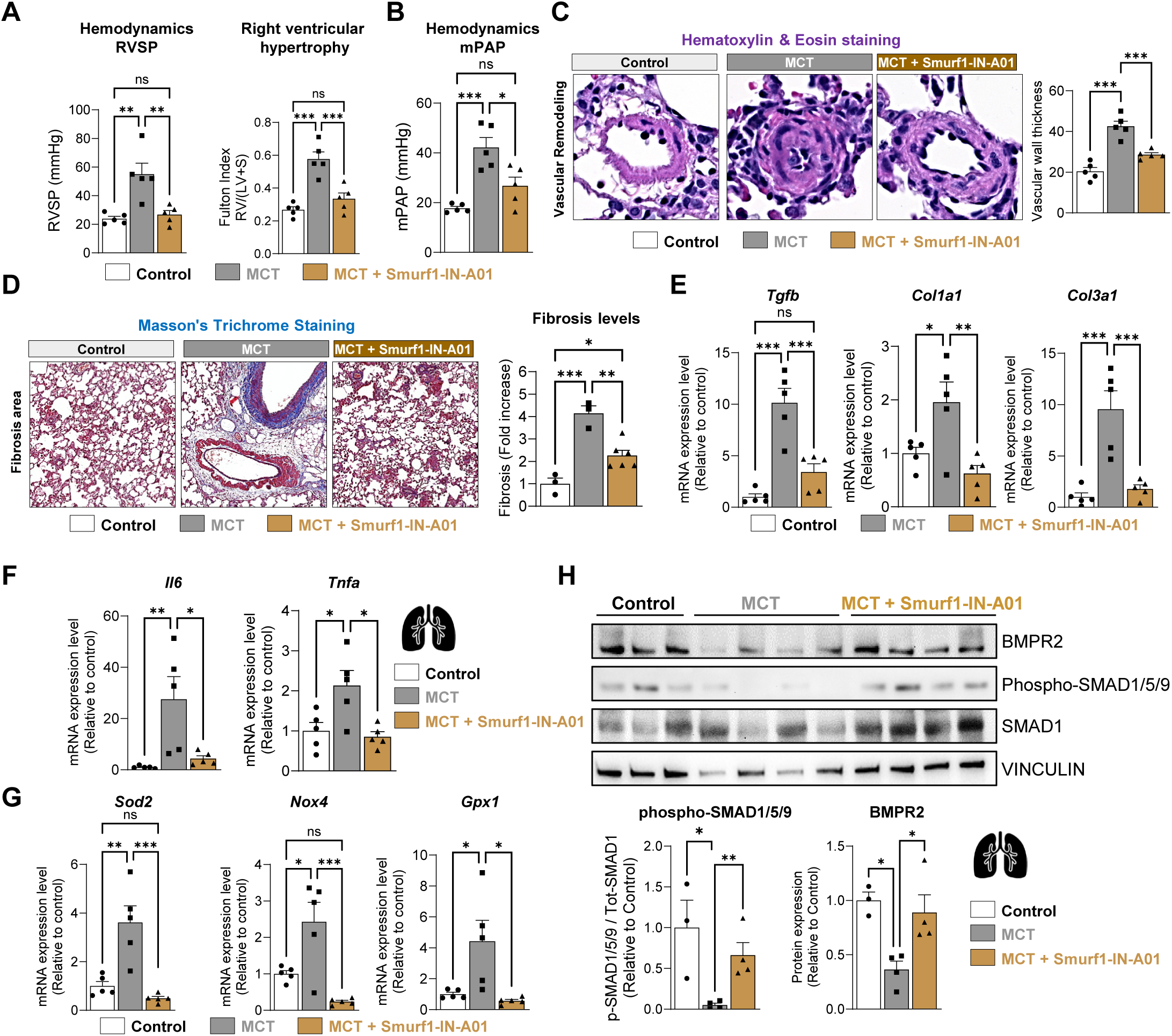
Smurf1 inhibition improves cardiopulmonary function and attenuates vascular remodeling in vivo. **(A)** Right heart catheterization in monocrotaline (MCT)-treated rats revealed reduced right ventricular systolic pressure (RVSP; left panel) and improved right ventricular hypertrophy, assessed by Fulton Index (right panel; ratio of RV to LV+S weights), following Smurf1-IN-A01 administration (n=5). **(B)** Mean pulmonary arterial pressure (mPAP) was measured via right heart catheterization in MCT rats post-treatment with Smurf1-IN-A01 (n=5). **(C)** Pulmonary vascular remodeling was quantified by hematoxylin and eosin (H&E) staining, assessing medial wall thickness of pulmonary arteries (n=5). **(D)** Masson’s trichrome staining visualized perivascular collagen deposition; quantification of fibrotic area was expressed relative to control animals (n=3-5). **(E–G)** Quantitative PCR analysis of lung tissues measuring mRNA expression of fibrotic markers (**E**, *Col1a1*, *Col3a1*, *Tgfb1*), inflammatory cytokines (**F**, *Il6, Tnfa*), and oxidative stress markers (**G**, *Nox4, Sod2, Gpx1*); n=5 per group. **(H)** Western blot analysis of lung lysates for BMPR2 protein expression and phospho-Smad1/5/9 signaling in Smurf1-IN-A01–treated rats (n=3-4). Data are presented as mean ± SEM. *p<0.05; **p<0.01, ***p<0.001.

## DISCUSSION

This study uncovers a pivotal epigenetic mechanism by which Smurf1, a HECT-type E3 ubiquitin ligase, disrupts BMPR2 signaling in PAECs, thereby driving endothelial dysfunction and vascular remodeling in PAH. Our results demonstrate that Smurf1 expression is markedly upregulated in PAECs derived from PAH patients and in experimental PAH models, consistent with prior reports by Murakami and colleagues, who first established Smurf1’s role in promoting BMPR2 and Smad1/5 ubiquitination and degradation ^19^. These findings align with subsequent work by Rothman et al., which confirmed Smurf1 enrichment in the pulmonary vasculature of PAH patients ^41^. Loss of BMPR2 signaling is a well-established hallmark of PAH, with mutations or downregulation of BMPR2 linked to poor clinical outcomes and exaggerated vascular remodeling^42^. Our findings extend this paradigm by identifying Smurf1 as a post-translational suppressor of BMPR2 and provide direct mechanistic evidence that pharmacological inhibition of Smurf1 restores BMPR2 levels and downstream Smad1/5/9 signaling in PAECs. This restoration correlates with reduced endothelial proliferation, consistent with the known antiproliferative role of BMPR2 in maintaining pulmonary vascular quiescence.

We further demonstrate that Smurf1 transcriptional upregulation is driven by epigenetic activation through H3K27 acetylation, mediated by the histone acetyltransferase EP300. These findings are corroborated by recent work from Chelladurai et al., who showed that global increases in H3K27ac contribute to aberrant activation of pathogenic transcriptional programs in PAH lungs, with EP300 emerging as a critical node linking inflammatory and proliferative gene expression ^38^. By integrating chromatin immunoprecipitation, transcriptomic, and pharmacologic analyses, we identify *SMURF1* as a novel transcriptional target of EP300, and we show that inhibition of EP300 using A485 reduces H3K27ac levels at the Smurf1 promoter, resulting in suppression of Smurf1 expression and restoration of BMPR2 signaling in PAH-PAECs. Our results underscore the significance of chromatin remodeling in modulating PAH-associated gene networks and suggest that targeting histone acetyltransferases like EP300 may normalize the expression of critical pathogenic mediators such as Smurf1. Building on our in vitro results, we employed the MCT-induced PAH rat model to test the therapeutic efficacy of Smurf1 inhibition. Treatment with the Smurf1-specific inhibitor A01 significantly reduced RVSP, attenuated pulmonary vascular remodeling, and suppressed fibrosis and inflammatory gene expression in lung tissue. However, these effects were only partially restorative, with disease parameters not fully normalizing to control levels. This partial rescue underscores the multifactorial nature of PAH. Further validation in additional preclinical models, such as the Sugen/hypoxia rats, which more accurately recapitulate obliterative vascular pathology, will be essential to evaluate the translational relevance and therapeutic durability of Smurf1-targeted interventions. Importantly, A01 restored BMPR2 protein expression and enhanced phosphorylation of downstream Smad1/5/9 signaling components, suggesting that Smurf1 inhibition rescues BMP pathway activity in vivo. Our findings align with a recent study demonstrating that Smurf1 inhibitors can reverse experimental PAH^43^, further positioning Smurf1 as a druggable effector whose modulation addresses a key mechanistic deficit in PAH pathogenesis.

Our findings support a dual-targeting approach that combines Smurf1 inhibition with BMPR2 gene augmentation to restore receptor abundance and limit its degradation simultaneously. This strategy may more effectively reconstitute BMP signaling fidelity and vascular homeostasis. Indeed, preliminary in vitro studies revealed that co-delivery of BMPR2 and Smurf1 inhibition synergistically enhanced SMAD1/5/9 activation and suppressed endothelial proliferation more robustly than either intervention alone. These data provide a mechanistic rationale for combining gene therapy with post-translational stabilization to overcome current therapeutic limitations. Further validation of this approach in additional models, such as the Sugen/hypoxia mouse, which recapitulates severe obliterative arteriopathy, will be essential to assess translatability, durability, and efficacy across PAH subtypes.

While our study identifies a novel epigenetic EP300-Smurf1 axis driving BMPR2 suppression and endothelial dysfunction in PAH, important questions remain. Specifically, the upstream signaling cues that govern EP300 activity in the pulmonary endothelium are not yet defined. Moreover, although our findings are supported by consistent trends across independent PH models, we acknowledge that expanding the sample size in future experiments would further strengthen the statistical power and generalizability of our results. In particular, including both male and female animals will be essential to account for potential sex-based differences in Smurf1 regulation and PAH pathobiology. Long-term studies will be necessary to assess the pharmacokinetics, safety, and therapeutic durability of Smurf1-targeted compounds in PH models. Furthermore, given the clinical heterogeneity of PAH, future investigations should explore the impact of sex, genetic background, and disease etiology on therapeutic response. Co-delivery strategies incorporating BMPR2 gene transfer with Smurf1 inhibitors represent a promising next step toward precision-guided therapy for PAH.

## CONCLUSION

This study identifies Smurf1 as a critical epigenetically regulated driver of BMPR2 degradation and endothelial dysfunction in PAH. Through integrative molecular and pharmacological analyses, we show that Smurf1 expression is governed by EP300-mediated H3K27 acetylation and that pharmacological inhibition of either Smurf1 or EP300 restores BMPR2 expression and reactivates downstream BMP signaling in pulmonary artery endothelial cells. In vivo, Smurf1 inhibition ameliorates pulmonary hemodynamics and vascular remodeling in a preclinical model of PAH. These findings highlight Smurf1 as a converging point between epigenetic dysregulation and impaired BMP signaling, two hallmarks of PAH pathogenesis, and establish its inhibition as a promising therapeutic strategy. Importantly, our preliminary combinatorial experiments suggest that pairing BMPR2 gene delivery with Smurf1 inhibition may synergistically enhance BMP pathway reactivation, offering a novel approach to overcome prior limitations of gene therapy alone. Taken together, our work provides mechanistic insight into the epigenetic control of Smurf1 and positions the Smurf1-BMPR2 axis as a therapeutically actionable pathway in PAH. Future studies should focus on advancing these findings toward clinical translation by refining dual-delivery platforms and evaluating long-term therapeutic efficacy in multiple PAH models.

## Ethics approval and consent to participate

This study utilized de-identified human lung tissue samples obtained from the Pulmonary Hypertension Breakthrough Initiative. Animal experiments were conducted in accordance with the National Institutes of Health Guide for the Care and Use of Laboratory Animals and were approved by the Institutional Animal Care and Use Committee (IACUC) of New York Medical College. Research involving recombinant or synthetic nucleic acid molecules and biohazardous agents was reviewed and approved by the Institutional Biosafety Committee (IBC) of New York Medical College.

## Consent for publication

This study utilized de-identified human lung tissue samples obtained from the Pulmonary Hypertension Breakthrough Initiative (PHBI). All samples were collected with informed consent under protocols approved by the respective Institutional Review Boards (IRBs) of participating institutions. As the data were de-identified and no individual-level information is presented, additional consent for publication is not required.

## Availability of data and materials

The datasets generated and analyzed during the current study are available from the corresponding author upon reasonable request.

## Authors’ contributions

MB conceived and designed the study, supervised the research, and wrote the manuscript. CL, MTO, and MB performed the experiments and contributed to data analysis. CL and MTO assisted with manuscript preparation and figure generation. JDE provided important conceptual insights and contributed to manuscript editing. All authors contributed to the editing of the manuscript and approved the final version.

## Acknowledgments

We thank the Pulmonary Hypertension Breakthrough Initiative (PHBI) for providing human lung biospecimens, and pulmonary artery endothelial cells from FD and PAH patients. We are grateful to the staff of the New York Medical College Animal Facility for their assistance with in vivo procedures. The authors also acknowledge the contributions of the Histopathology Core for tissue processing and staining.

