## Supplementary Figure 1 for "Epigenetic Activation of Endothelial Smurf1 via EP300-Mediated H3K27ac Disrupts BMPR2 Signaling in Pulmonary Arterial Hypertension"

**Supplementary figure legends**

**Supplementary Figure 1. SMURF1 promotes the expression of pro-inflammatory cytokines in PAEC.** **A)** Quantitative PCR analysis of *IL6* (left panel) and *TNFA* (right panel) mRNA levels in PAH-PAECs following shRNA-mediated knockdown of *SMURF1* (shSMURF1) compared to non-silencing control (shNS), n = 4. **B)** *SMURF1* transcript (left panel) and protein levels (right panel) in donor-derived PAECs (FD-PAECs) transfected with a *SMURF1* expression construct or empty vector control (pCMV.SPORT6). **C)** QPCR quantification of *IL6* (left panel) and *TNFA* (right panel) mRNA expression in FD-PAECs overexpressing *SMURF1* or empty vector control. Data are presented as mean ± SEM. $^{**}p < 0.01$; $^{***}p < 0.001$.

### Supplementary Figure 1

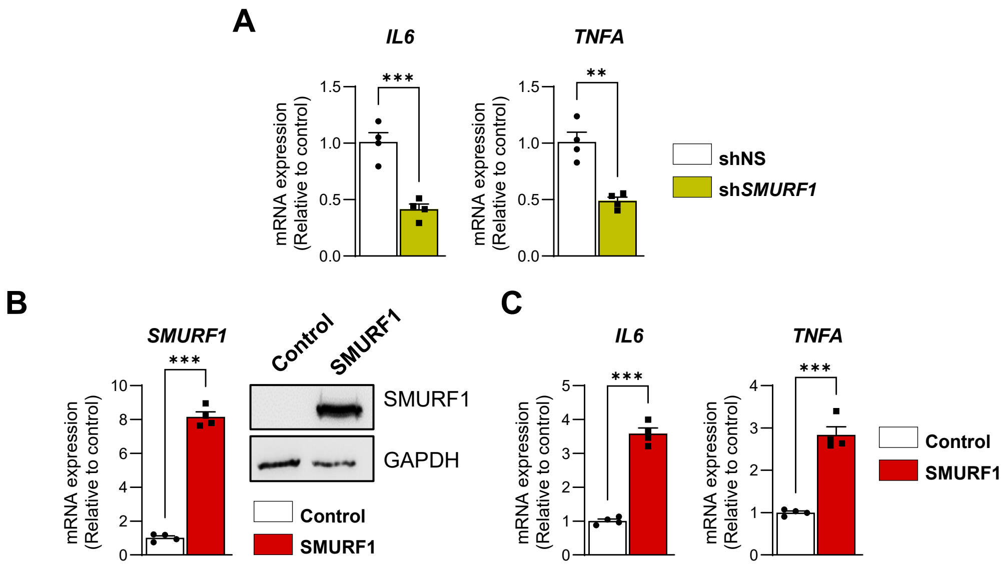
