## Supplemental Table 1 for "Epigenetic Activation of Endothelial Smurf1 via EP300-Mediated H3K27ac Disrupts BMPR2 Signaling in Pulmonary Arterial Hypertension"

**Supplementary Table 1.** Primer sequences for RT-qPCR and ChIP-qPCR analysis; clone ID and catalog numbers for shRNAs (Open Biosystems); antibodies used; source, and concentration of chemical inhibitors used.

| **Application** | **Gene symbol** | **Species** | **Forward primer (5′-3′)** | **Reverse primer (5′-3′)** |
|---|---|---|---|---|
| **RT-qPCR** | SMURF1 | Mouse Rat Human | TCAACCGACACTGTGAAAAACA | TCTGCTGATGGCATTGGAGAG |
| | COL1A1 | Rat | TTCAGTGGTTTGGATGGTGCCAA | CCAGCTTCACCCTTAGCACCA |
| | COL3A1 | Rat | GAGATGTCTGGAAGCCAGAACCATG | ATCTCCCTTGGGGCCTTGAGGT |
| | TGFB | Rat | CCTGCAAGACCATCGACATGGAG | GGTCGCGGGTGCTGTTGTA |
| | ANP | Rat | CCCGACCCACGCCAGCATGG | CAACTGCTTTCTGAAAGGGGT |
| | BNP | Rat | ACAATCCACGATGCAGAAGCT | GGGCCTTGGTCCTTTGAGA |
| | β-MHC | Rat | ACAGAGGAAGACAGGAAGAACCTAC | CACAAGATCTACTCCTCATTCAG |
| | SOD2 | Rat | AACGCGCAGATCATGCAGCT | TTCAGTGCAGGCTGAAGAGC |
| | NOX4 | Rat | TGGATGACTGGAAACCATACAAG | GACTGAGGTACAGCTGGATG |
| | GPX1 | Rat | GGTGCTGCTCATTGAGAATGTC | TCTTGCCATTCTCCTGATGTCC |
| | IL-8 | Human | TTGCCAAGGAGTGCTAAAGAACTTA | AGCTCTCTTCCATCAGAAAGCTTTA |
| | TNFα | Human | CTTGTTCCTCAGCCTCTTCTCCTTC | GGGTTCGAGAAGATGATCTGACTG |
| | | Rat | ATCCGAGATGTGGAACTGGC | CGATCACCCCGAAGTTCAGT |
| | IL1β | Human | ATGATGGCTTATTACAGTGGCAATG | ATCTTCCTCAGCTTGTCCATGG |
| | IL6 | Human | AGCTGCAGGCACAGAACCAG | ATTTGCCGAAGAGCCCTCAG |
| | | Rat | GGTCTTCTGGAGTTCCGTTTCT | AGAGCATTGGAAGTTGGGGTAG |
| | 18S | Mouse Rat Human | GTAACCCGTTGAACCCCATT | CCATCCAATCGGTAGTAGCG |
| | EP300 | Human | CTAATTCTCTCTCCAATCAAGTGCG | TGGCAAAATGCCCTTGTTCCAT |
| **ChIP-qPCR** | SMURF1 Promoter | Human | GAAGCAGCGAGGGTCAGAG | GAGCCGGGACACAAACTCC |
| **shRNAs** | **Gene symbol** | **Clone ID** | | **Catalog number** |
| | *SMURF1* | TRCN0000003473 | | RHS3979-201735746 |
| **Overexpression** | **Gene symbol** | **Clone ID** | | **Catalog number** |
| | *BMPR2* | ccsbBroad304_00169 | | OHS6085-213572865 |
| **Immunoblotting** | **Protein symbol** | **Antibody source** | | **Dilution** |
| | SMURF1 | Cell signaling | | 1:1000 |
| | H3K27ac | Cell signaling | | 1:1000 |
| | Histone H3 | Cell signaling | | 1:1000 |
| | Phospho-SMAD1/5/9 | Cell signaling | | 1:1000 |
| | Total SMAD1 | Cell signaling | | 1:1000 |
| | BMPR2 | Invitrogen | | 1:1000 |
| | VINCULIN | Cell signaling | | 1:5000 |
| | GAPDH | Invitrogen | | 1:5000 |
| Pharmacological agents | **Compounds** | **Concentration** | | **Source** |
| | A485 | 2 μM | | MedChem Express |
| | Smurf1-IN-A01 | 20 μM | | MedChem Express |
